# The spectral sensitivity of *Drosophila* photoreceptors

**DOI:** 10.1101/2020.04.03.024638

**Authors:** Camilla R. Sharkey, Jorge Blanco, Maya M. Leibowitz, Daniel Pinto-Benito, Trevor J. Wardill

## Abstract

*Drosophila melanogaster* has long been a popular model insect species, due in large part to the availability of genetic tools and is fast becoming the model for insect colour vision. Key to understanding colour reception in *Drosophila* is in-depth knowledge of spectral inputs and downstream neural processing. While recent studies have sparked renewed interest in colour processing in *Drosophila*, photoreceptor spectral sensitivity measurements have yet to be carried out *in vivo*. We have fully characterised the spectral input to the motion and colour vision pathways, and directly measured the effects of spectral modulating factors, screening pigment density and carotenoid-based ocular pigments. All receptor sensitivities had significant shifts in spectral sensitivity compared to previous measurements. Notably, the spectral range of the Rh6 visual pigment is substantially broadened and its peak sensitivity is shifted by 92 nm from 508 to 600 nm. We propose that this deviation can be explained by transmission of long wavelengths through the red screening pigment and by the presence of the blue-absorbing filter in the R7y receptors. Further, we tested direct interactions between photoreceptors and found evidence of interactions between inner and outer receptors, in agreement with previous findings of cross-modulation between receptor outputs in the lamina.

## Introduction

Colour cues in the natural environment are used by many insects, guiding behaviours such as prey detection, mate selection and more general tasks such as flight navigation. To detect and distinguish wavelength cues, the output of two or more photoreceptors with different spectral sensitivities must be compared. This colour-opponency system has been well explored in vertebrates but only more recently have insect colour opponent neurons been characterised, using *Drosophila* as the model^1,2^. *Drosophila melanogaster* is fast becoming the model for insect colour vision due to the wide range of genetic tools available. Surprisingly, despite new advances in our understanding of more complex motion and colour visual processing in *Drosophila*, the characterisation of spectral sensitivity in this species has not been investigated since it was first revealed 20 years ago^3,4^.

The *Drosophila* compound eye is formed from approximately 800 ommatidial units, each comprising 6 outer (R1 - 6) and 2 inner photoreceptors (R7 and R8) with an open rhabdom structure (**Fig. 1a**). Sensitivity of the photoreceptors is largely determined by the underlying visual pigment and ommatidia can be subdivided into two major classes, ‘pale’ (p) or ‘yellow’ (y), owing to their appearance under the microscope, with the latter possessing a blue-absorbing yellow filter in the R7y receptor alongside the UV-sensitive Rh4 visual pigment^5^. In the outer receptors of all ommatidia, opsin Rh1 confers broadband blue-green sensitivity (478 nm) with an additional peak in the UV due to an associated carotenoid-derived sensitising pigment^4,6^. Inner receptor R7 cells express UV-sensitive Rh3 (R7p) or Rh4 (R7y) and the proximal receptor R8 cells express either blue-sensitive Rh5 (R8p) or green-sensitive Rh6 (R8y)^3,4^. Direct intracellular recordings of the inner photoreceptors have not been possible due to their small size and stochastic distribution across the retina. Instead, peak sensitivities for Rh3 – Rh6 have been estimated from microspectrophotometery (MSP), visual pigment extracts and electroretinography (ERG) with ectopic expression of inner receptor opsin in the more numerous outer receptors, using white-eyed *Drosophila*^3,4^. Although this enabled the underlying visual pigment sensitivities to be measured (Rh3 – Rh6: 345, 375, 437, 508 nm; **Fig. 1b**), these studies were unable to quantify the sensitivity of each when measured *in vivo*, in the photoreceptor cells where the opsins are normally expressed along with their naturally associated ocular screening pigment and photoreceptor filtering pigments (**Fig. 1 c, d and e)**.

**Figure 1.**
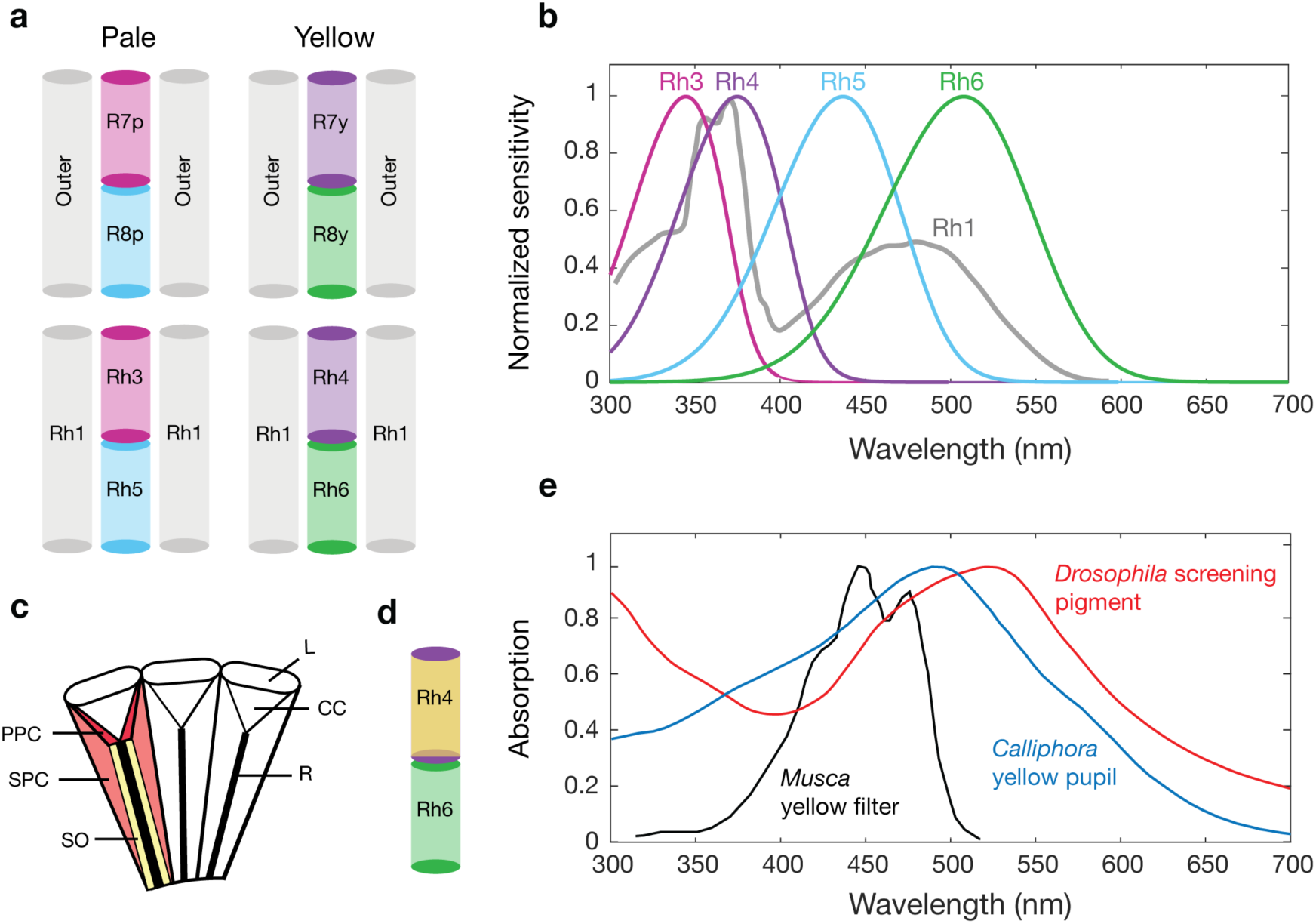
An overview of Drosophila photoreceptors, visual pigments and fly ocular pigments. (a) The arrangement of inner R7/R8 and outer receptors in pale- and yellow-type ommatidia of Drosophila and the opsins expressed in each. (b) Spectral sensitivities of visual pigments Rh3 – 6 modelled using visual pigment templates and previous sensitivity estimates^4^. Rh1 has a characteristic shape owing to a blue-green sensitive visual pigment coupled to a UV-sensitising pigment. (c) Longitudinal section diagram of an insect compound eye indicating the distribution of screening pigment in primary pigment cells (PPC) and secondary pigment cells (SPC) that optically isolate the corneal lens (L), crystalline cone (CC) and rhabdom (R). The soma (SO) of the photoreceptors contain mobile pigment granules that form the fly pupil. (d) Location of the blue-absorbing yellow filter alongside opsin Rh4 in the R7y rhabdom. (e) Absorption of red Drosophila screening pigment, Calliphora yellow pupillary pigment and the blue-absorbing yellow filter measured in Musca^5,16,28^.

The spectral sensitivity of a cell is not dictated solely by the underlying visual pigment but is shaped also by numerous optical filters, including screening pigment that optically isolates neighbouring ommatidia and is located in the primary and secondary pigment cells (**Fig. 1c**). Pupillary pigment is also present in the soma of the photoreceptors and modulates light input to the rhabdom. In addition to this, dipteran flies have carotenoid-based pigments: a sensitising pigment that contributes additional sensitivity in the UV through energy transfer to the visual pigment (Rh1; **Fig. 1c**) and a blue-absorbing yellow filter in the R7y cells^5^ (**Fig 1.d and e**). *Drosophila* screening pigment absorbance peaks at 525 nm and becomes more transmissive at longer wavelengths (**Fig. 1e**), an adaptation thought to maximising the reconversion of Rh1 rhodopsin from its metarhodopsin, which peaks at 566 nm^4,7^. Early studies pointed towards a red receptor in dipterans^8^, later shown to be an artefact of long wavelength light leakage^9^. It has been argued that this red sensitivity may be of little consequence to the visual system of such flies when under ecologically-relevant levels of illumination ^9^. Furthermore, it is thought that the wavelength peak of *Drosophila* Rh6 (508 nm) is short-wavelength shifted when compared to flies with brown screening pigment (e.g. *Musca*, 520 nm), as an adaptation to reduce the absorption of red light. The effect of light leakage on the *Drosophila* Rh6 visual pigment has yet to be tested *in vivo*.

Much of what is understood about animal colour vision is derived from studies of vertebrates and only more recently have investigations revealed the neural basis of colour information processing in insects. Colour information processing in *Drosophila* has been investigated using behavioural experiments^10,11^, modelling^12^ and by visualising neural responses directly in the fly brain, in response to light stimulation with genetic manipulation^1,2^. Colour opponency begins at the first visual synapse, the photoreceptor terminal, with reciprocal inhibition occurring between paired R7 and R8 receptors^1^. Additionally, there is transfer of information between inner and outer receptors *via* gap junctions in the lamina that is thought to enhance the sensitivity of the motion detection pathway^13^. However, it is not clear yet whether interactions do occur directly between inner photoreceptors, as has been demonstrated in other insects^14,15^ and whether opponency can be detected at the photoreceptor level.

Here we report the first complete *in vivo* characterisation of *Drosophila* spectral sensitivity, for each receptor type, by selectively restoring photoreceptor activity in flies with no receptor activity (*norpA*) and testing response using ERG. We characterise the sensitivity of photoreceptors flies with screening from distal receptors and ocular pigments intact and we test the effect of screening pigment and the blue-absorbing yellow filter on inner receptors. We reveal significant shifts in spectral sensitivity for all receptor sensitivities and a large 92 nm shift in sensitivity of the Rh6 visual pigment when measured in its native photoreceptor (R8y) from 508 nm to 600 nm. We also find that the blue-absorbing yellow filter refines sensitivity of the Rh4 visual pigment in R7y photoreceptors. Furthermore, we explore the effect of reciprocal inhibition between inner photoreceptors on spectral response at the level of the photoreceptors and to what extent the outer photoreceptors input to this system. Our results indicate that the input from inner photoreceptors to downstream neuronal process is linear and provides no clear evidence for direct interactions at the photoreceptor level. Spectral modulation can be seen however between inner and outer receptors, providing further evidence of interactions between motion and spectral channels in the lamina.

## Results

### Single opsin rescues, Rh1, Rh3, Rh4, and Rh5

We generated flies with rescued activity of specific photoreceptor types (single opsin rescue flies) to test the spectral sensitivity of the inner R7p (Rh3), R8p (Rh5), R7y (Rh4), R8y (Rh6) and outer receptors (Rh1), using ERG. We were able to measure reliably from all single opsin rescue genotypes. Rh1 rescue flies exhibited characteristic on and off transients at the start and end of the 200 ms light pulse, absent in Rh3 – Rh6 (**Fig. 2**) rescue flies. The spectral responses of rescue flies Rh3, Rh4 and Rh5 with wild-type screening pigment had significant shifts in spectral sensitivity for portions of their detection range compared with previous characterisations (**Fig. 3**, black dashed traces). Sensitivities of Rh3 and Rh4 rescue flies were significantly short wave shifted from previous estimates, peaking at 330 and 355 nm, respectively (previous estimates: 345 and 375 nm). The peak sensitivity of the Rh5 rescue flies (435 nm) was similar compared to previous measurements of the visual pigment (437 nm^4^) but the response had significantly boosted sensitivity in the ultraviolet range. The spectral sensitivity response of R1 - R6 (Rh1) was also significantly altered from a simple visual template. The red end of the spectrum was depressed and there were notably three fluctuations in the waveform around the peak (**Fig. 3**). Spectral curves from all four photoreceptor cell types were broader than predicted by a visual pigment template at peak sensitivity.

**Figure 2.**
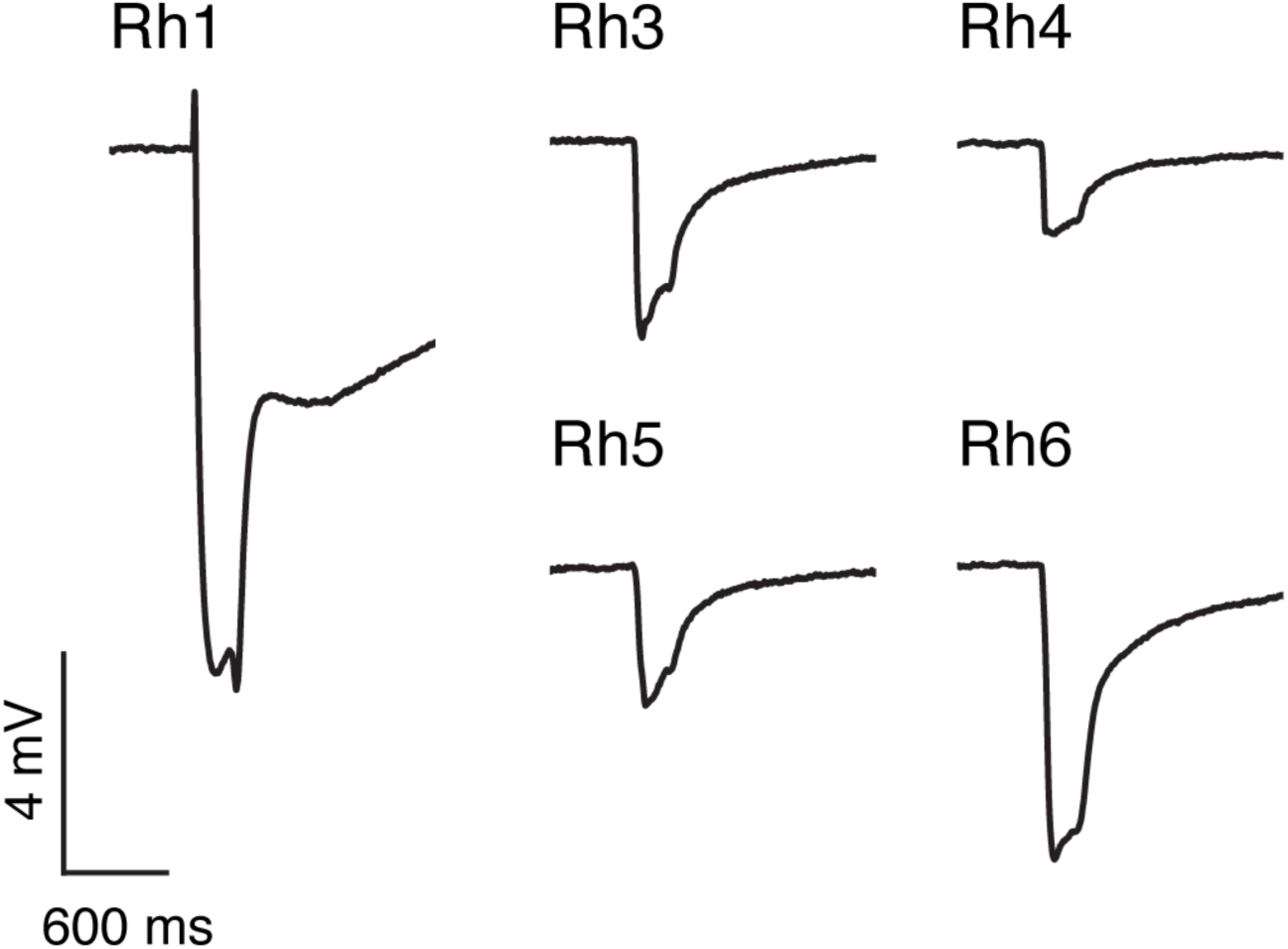
Example ERG traces of single opsin rescue flies in response to a 200 ms pulse of light at photoreceptor saturation (3.60 × 10^12^ photons^-1^m^-2^s^-1^).

**Figure 3.**
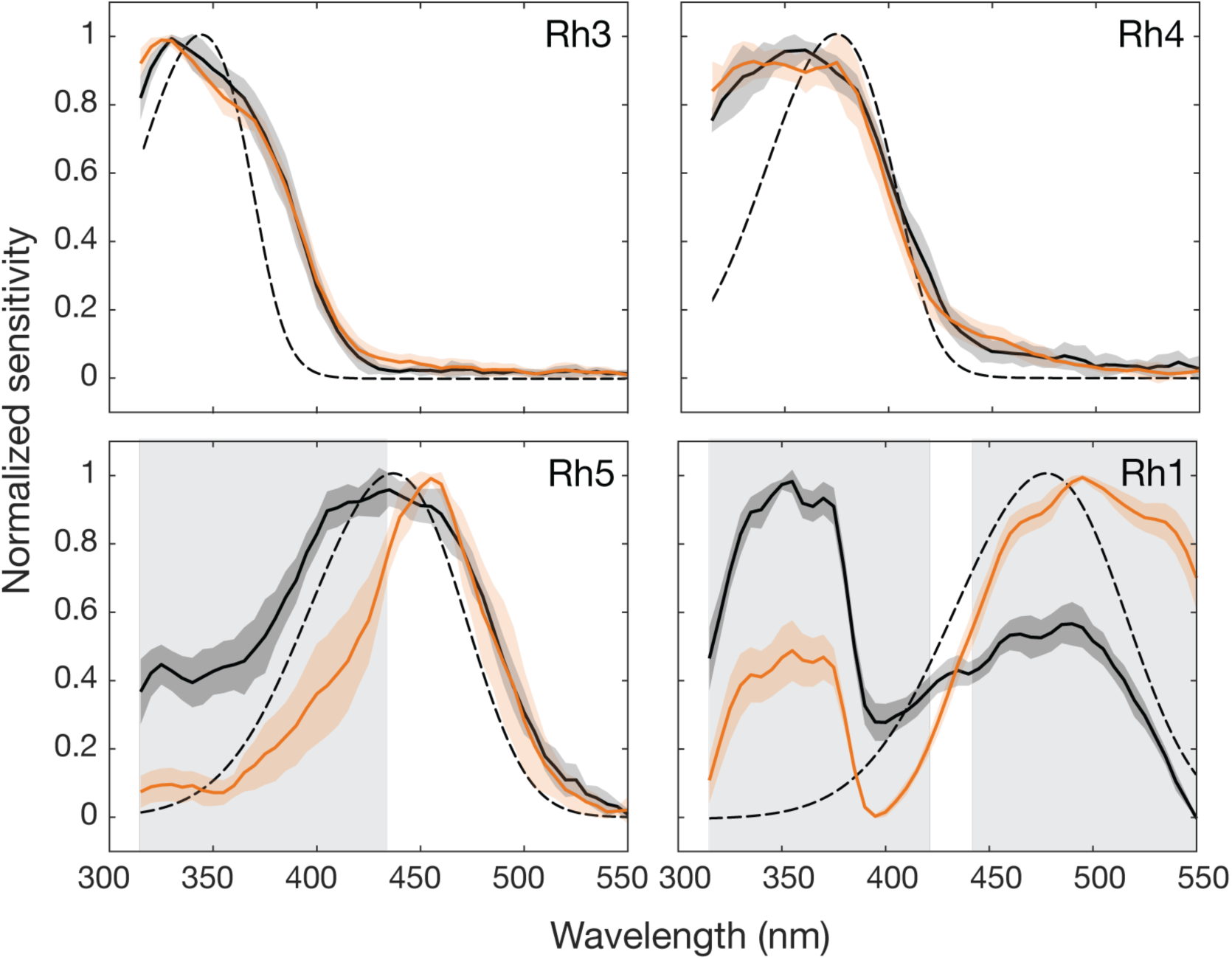
Spectral sensitivity of red and orange eye flies with selectively rescued photoreceptor responses. Normalized spectral sensitivity of red eye (black lines) and orange eye (orange lines) flies with rescued activity of Rh1, Rh3, Rh4 or Rh5. Modelled visual pigment templates (dashed lines), based on previous estimates ^3,4,29^. Error shown is standard deviation. Shading denotes significance between red and orange eye flies using a two-sample Student’s t-test at p<0.001. For Vlog-I curves, see Supplementary Figure (**S1**).

To test the effect of screening pigment on photoreceptor response, we examined opsin rescue flies with wild-type (red-eye) and reduced screening pigment (orange-eye). The spectral sensitivities of Rh3 and Rh4 single opsin rescue flies were not affected by a reduction of screening pigment. However, a reduction in screening pigment led to narrowing of the Rh5 rescue response (**Fig. 3**, orange traces) and a bathochromic (long-wave) shift in the peak sensitivity by 20 nm, from 435 nm to 455 nm. The spectral profile of Rh1 rescue flies shows the characteristic triple-peaked UV spectrum of the UV-sensitising pigment coupled with curve of the visual pigment peaking at 485 – 490 nm (**Fig. 3**, orange traces). Screening pigment reduction caused a shift in sensitivity towards the visual pigment peak, reducing the relative sensitivity in the UV region.

Dietary carotenoids were removed from the diet of red eye rescue flies to test for the presence of carotenoid pigments in the outer receptors (Rh1) and inner R7p (Rh3), R7y (Rh4) and R8p (Rh5) receptors. The response of the UV-sensitising pigment coupled to visual pigment Rh1 was effectively removed by carotenoid deprivation after one generation on yeast-glucose food and showed no further change in spectral shape after two generations (**Fig. 4**). Carotenoid deprivation had no effect on the spectral response of Rh3 opsin rescue flies but broadened the response in Rh4 rescue flies above 400 nm (**Fig. 4**). Responses were absent in Rh5 rescue flies when carotenoids were removed from the diet.

**Figure 4.**
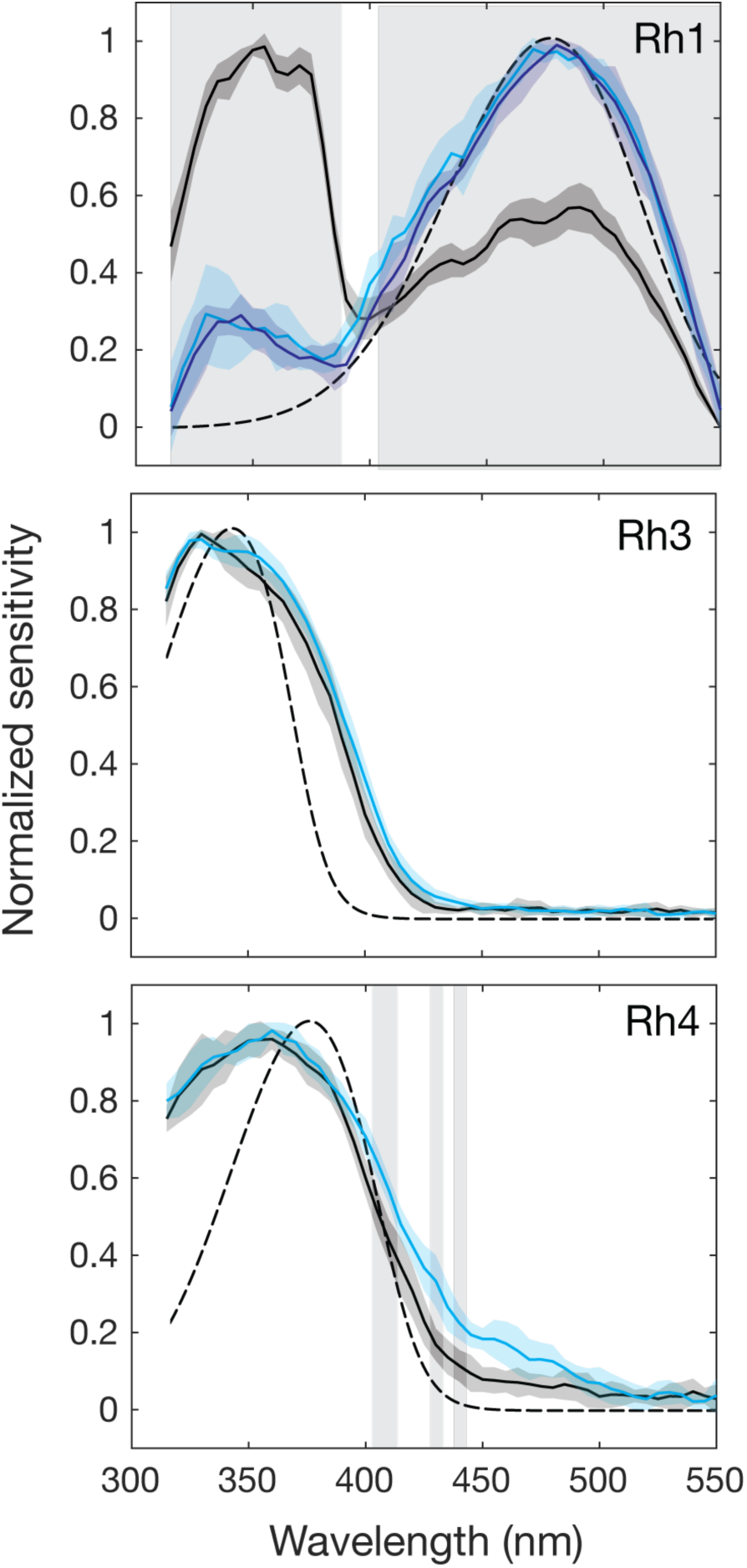
Spectral sensitivity of flies with carotenoid deprivation and selectively rescued photoreceptor responses. Normalized spectral sensitivity of red-eye flies raised on a regular diet of yellow cornmeal (black) or carotenoid-deprived flies raised on yeast-glucose for one (light blue) or two (dark blue) generations, in the case of Rh1 flies. Modelled visual pigment templates (dashed lines) based on previous estimates^3,4,29^. Error shown is standard deviation. Shading denotes significance between normal and carotenoid-deprived red-eye flies using a two-sample Student’s t-test at p<0.001.

### Single opsin rescue Rh6

In Rh6 single opsin rescue flies, where R8y activity was rescued, the spectral response was far broader than expected and considerably shifted in peak wavelength sensitivity from the previous estimate (508 nm) to 600 nm when the sensitivity was measured *in vivo* (**Fig. 5a**). This large bathochromic shift was reversed by 45 nm to 555 nm by the reduction of screening pigment in orange-eye flies. Sensitivity of white-eye mutants where screening pigment was absent and Rh6 was expressed in the outer receptors peaked close to the predicted peak wavelength of the visual pigment (508 nm), at 510 nm (**Fig. 5a**).

**Figure 5.**
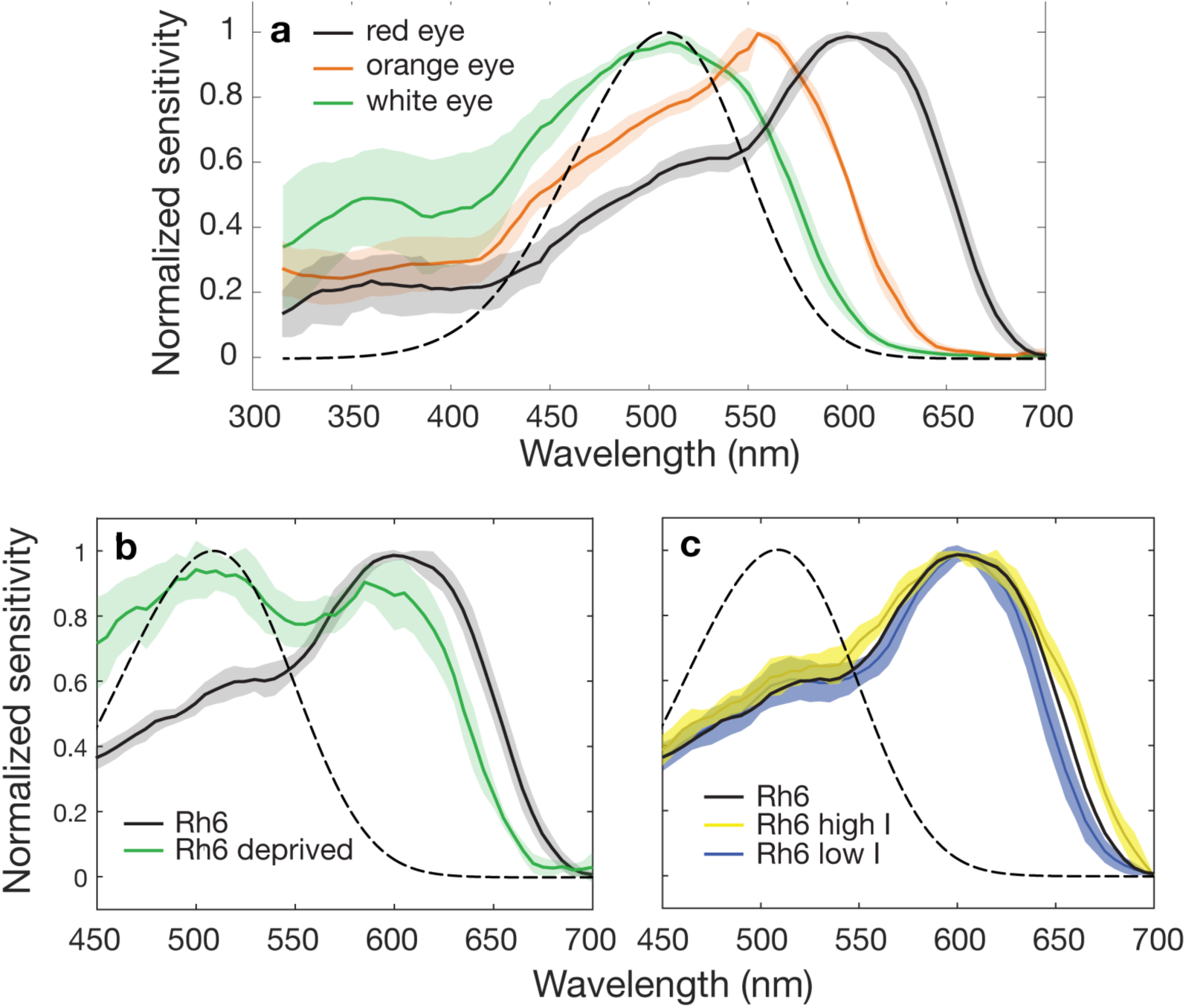
The effect of screening pigment, carotenoid deprivation and ectopic expression on the spectral sensitivity of the Rh6 visual pigment. (a) Normalized spectral sensitivity of Rh6 rescue flies with red- (black solid), orange- (orange) or white eyes with Rh6 expressed in outer receptors (green), peaking at 600, 555 and 510 nm, respectively. Spectral sensitivity template of the Rh6 visual pigment peaking at 508 nm (dashed line). (b) Normalised spectral sensitivity of Rh6 rescue flies with red eyes raised on a regular (black solid line) or carotenoid-free diet for two generations (green line). (c) Response curves of Rh6 rescue flies tested 0.5 log units above (high I) and below (low I) normal test intensity. All error shown is standard deviation.

We aimed to test the effect of the carotenoid-based blue-absorbing yellow pigment on the sensitivity of Rh6 by means of carotenoid deprivation. When the yellow pigment was removed, two peaks of sensitivity could be seen, one at the peak sensitivity of Rh6 (508 nm) and the other close to the peak of the flies raised on regular carotenoid rich diet (600 nm) (**Fig. 5b**). There was no change in the shape of the spectral response between one and two generations of carotenoid deprivation (Supplementary Figure **S2**). As carotenoid deprivation reduces the overall sensitivity of the photoreceptor by chromophore depletion, flies must be tested at a higher light intensity. As such, due to the limit of light in the system, flies were tested at the lower end of the VlogI curve. To simulate these potential confounding conditions, we tested Rh6 rescue flies raised on a regular diet, at 0.5 log units of light above and below the normal testing intensity. Spectral responses were relatively unchanged by light intensity with minor broadening and narrowing of the spectral curve occurring at higher and lower light intensities, respectively (**Fig. 5c**). Importantly however, the spectral shape of carotenoid-deprived Rh6 rescue flies could not be replicated.

### Double opsin rescues

To test for potential spectral modulation at the photoreceptor level, responses from flies with two active photoreceptor types (double opsin rescue flies) were compared with the sum of corresponding single photoreceptors. We found no difference in spectral shape when doubles were tested at different intensities, according to the two VlogI tests carried out at each visual pigment peak sensitivity, with the exception of a single wavelength in the Rh3 and Rh5 double rescue flies (Supplementary Figure **S3**). If there were interactions between photoreceptors we would expect the sum of the single receptors to differ from that of the double rescue. For R7 and R8 receptor pairs both in their corresponding ommatidial pairs (e.g. Rh3 and Rh5: R7p and R8p) and in non-corresponding ommatidial pairs (e.g. Rh4 and Rh6: R7p and R8y) showed highly similar responses to the sum of the single rescues with the exception of the Rh3 and Rh6 double rescue (**Fig. 5a**). Although a clear difference in spectral profile can be seen between the expected sum of Rh3 and Rh6 and the corresponding double opsin rescue (**Fig. 6a**), this difference is no longer present in the non-normalised data (Supplementary Figure **S4**). The sum of the Rh3 and Rh6 spectral responses do closely match those of the double rescue flies (Rh3 and Rh6) but only when tested at the intensity calculated from the VlogI experiment at the corresponding peak (Supplementary Figure **S4**).

**Figure 6.**
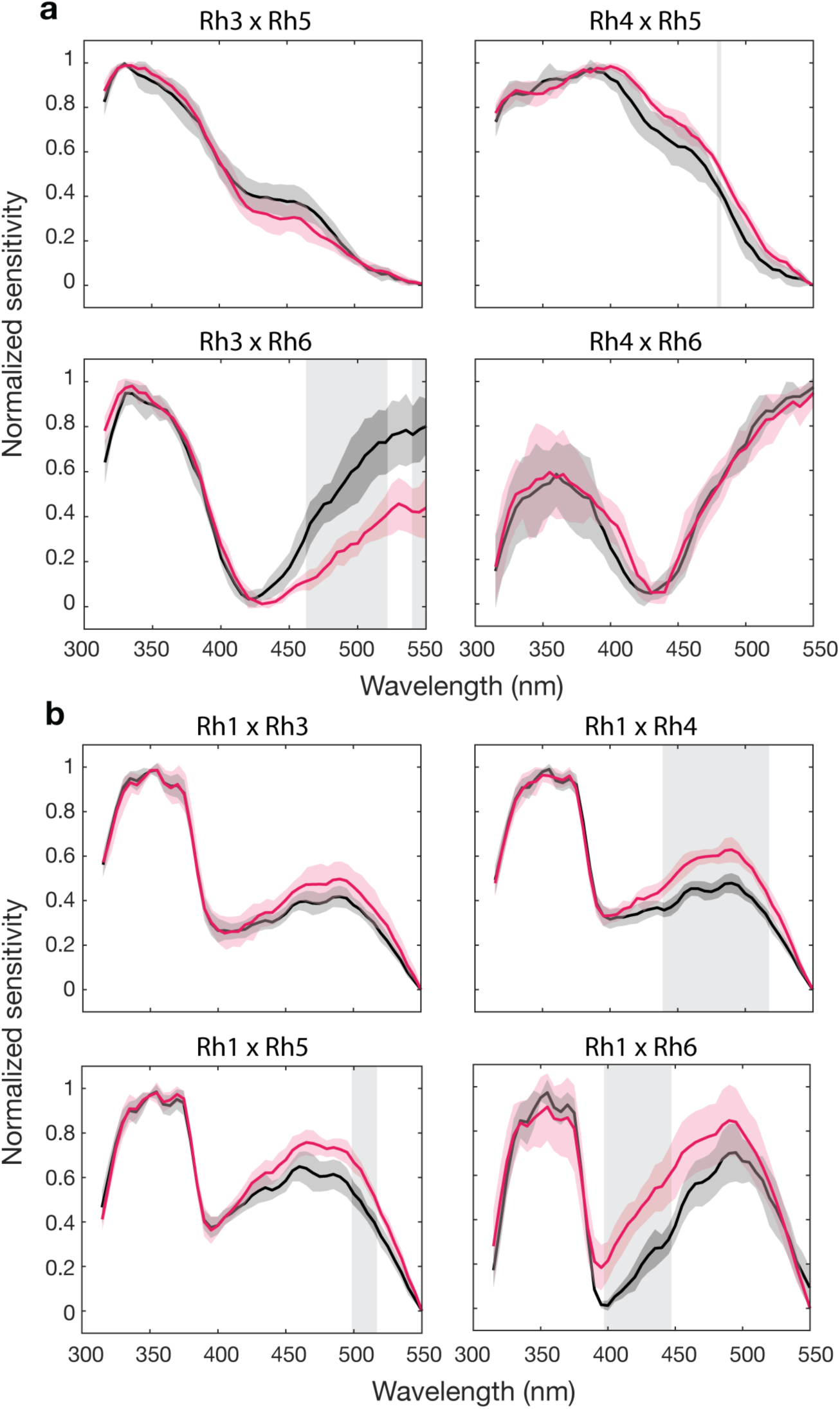
Spectral sensitivity of double opsin rescue flies and the sum of equivalent single rescue responses. (a) Normalized responses from double opsin rescue flies with the activity of two photoreceptor types active (pink) compared with the algebraic sum of the single rescue responses (black). Animals were tested at two intensities, derived from the VlogI response at Rh3/Rh4 peak sensitivities and Rh5/Rh6 peak sensitivities (not shown, see Supplementary Figure **S3**). (b) Double opsin rescue flies with Rh3 – 6 and Rh1 (pink) compared to the sum of single rescue flies (black). Double rescue flies were tested at intensities using VlogI responses at the Rh1 peak sensitivity and Rh3/Rh4/Rh5/Rh6 peak sensitivities (not shown, see Supplementary Figure **S3**). All error shown is standard deviation. Shading denotes significance between double rescue response and sum of singles using a two-sample Student’s t-test at p<0.001.

Significant differences in spectral shape between the expected sum of single photoreceptor responses and double opsin rescues between inner and outer receptors were observed for outer receptors in combination with Rh4, Rh5 and Rh6 (**Fig. 6b**). In all three cases, the observed shift in the sensitivity curve is due to a lower than expected response in the UV, which translates to a relatively higher response at wavelengths higher than 400 nm, upon normalization (Supplementary Figure **S4**). The sum of all mean responses from each photoreceptor type matches well to the response measured from wild-type flies (Supplementary Figure **S5**), indicating that the photoreceptor responses do indeed sum linearly overall.

## Discussion

We have shown that using norpA flies with activity rescued in selected photoreceptor types alongside ERG enabled the characterisation of spectral sensitivity in *D. melanogaster* as an alternative to intracellular recordings. The responses of inner R7 receptors are similar to their respective visual pigments Rh3 and Rh4^4^ but have significant changes induced by ocular screening and filtering pigments. The spectral profiles of R8 inner receptors are even more notably modified by the presence of ocular pigments and are poorly described by the underlying visual pigment sensitivities of Rh5 and Rh6^4^. The shift in sensitivity of Rh5 rescue flies when screening pigment levels were reduced is likely due to the increase in off-axis light, which increases direct stimulation of the R8p receptor. The resulting sensitivity curve more closely resembles the underlying sensitivity of the visual pigment^4^. This indicates that the broadening of sensitivity that we observe is likely due to distal screening from the UV-receptor R7p.

The sensitivity curve of Rh6 rescue flies (R8y) with wild-type (red) screening pigment is far broader than expected, with low levels of sensitivity in the UV and a major peak of sensitivity at 600 nm (**Fig. 5a**). This long-wavelength shift of 92 nm from 508 to 600 nm can be in part explained by the absorption curve of the red screening pigment, which is maximal at 290 and 525 nm^16^ (**Fig. 1e**). Above 525 nm there is a steady decline in light absorption by the screening pigment. This has the effect of increasing the light available to the R8y receptors above 525 nm, where it is still able to stimulate the long-wavelength tail of the Rh6 visual pigment. This effect can be seen by the hypsochromic (short-wave) shift in peak sensitivity of the reduced screening pigment mutants back towards the peak sensitivity of the Rh6 visual pigment to 555 nm (**Fig. 5a**). By reducing the effect of the long-wavelength light leakage, the relative absorption of the Rh6 is shifted towards the peak sensitivity of the underlying visual pigment. We are confident that the peak sensitivity of Rh6 in *Drosophila* is indeed close to the previously measured 508 nm as confirmed by our measurements of white eye flies with ectopic expression of Rh6 in the outer receptors (**Fig. 5a**). However, there is a clear effect of light leakage on the sensitivity of the R8y receptor that shifts sensitivity towards the red.

It has been proposed that the Rh1 and Rh6 of *Drosophila* and other red-eyed flies may be sensitive to the longer wavelengths of light that leak through the red screening pigment and as a consequence this would degrade spatial resolution^17^. At high light levels, the pupil response causes increased absorption of blue-green wavelengths, reducing the stimulation of the Rh1 visual pigment, instead favouring UV-sensitivity and photoconversion of metarhodopsin^18^. This may reduce absorption of stray long-wavelength light by the Rh1 visual pigment but no such mechanism is present for Rh6. Interestingly, while it is assumed that stray red light will negatively affect the spatial resolution of fly vision at long wavelengths, the relative absorption of the red screening pigment is near equal at both 400 and 600 nm. This suggests that the effects of off-axis light stimulation and scatter within the eye would serve to degrade spatial resolution at both wavelengths similarly.

Like other large dipterans, *Drosophila* has a blue absorbing carotenoid present in the distal R7y retinula cells^5^. In *Musca* and *Calliphora* this serves to reduce the light absorbed by the blue-sensitive visual pigment in R7y, instead conferring sensitivity to the UV by the presence of a UV-sensitising pigment^19,20^. This interesting and complex system is not found in *Drosophila*, rather UV-sensitivity is achieved simply by the presence of a dedicated UV-sensitive visual pigment Rh4^3^. However, the presence of this carotenoid pigment in *Drosophila* was not previously understood and its effect on R8y has not been tested until now. When carotenoids were removed from the diet of flies with rescued R8y activity, sensitivity was restored close to the peak of the visual pigment (510 nm) but some effect of light leakage remained at 600 nm. This would suggest that the blue-absorbing filter reduces light available to Rh6 at its peak sensitivity (508 nm) and instead extends sensitivity towards the red. These findings could not be replicated by simulating the differences in experimental conditions, either reducing or increasing testing intensity. One generation was sufficient to remove the contribution of the Rh1 UV-sensitising pigment and further generations of carotenoid deprivation did not change the response curve of Rh6 flies (Supplementary Figure **S2**), indicating that any carotenoid filters had been fully removed by one generation.

The effect of the carotenoid filter can also be seen in the Rh4 rescue flies, where it is located. When removed, sensitivity is increased at wavelengths greater than 400 nm suggesting that this filter narrows sensitivity in the UV, potentially increasing wavelength discrimination in that region of the spectrum. We suggest that this filter both contributes to refining the sensitivity of the UV-sensitive R7y (Rh4) photoreceptor and bathochromically shifting the sensitivity of R8y cells by depleting wavelengths of light between 400 and 540 nm in the R7y receptor. In large flies, the yellow filter only shifts R8y sensitivity from 520 to 540 nm but it is not known whether the filter in *Drosophila* absorbs at longer wavelengths, which could explain the larger shift we observe. Unfortunately the absorption properties of this carotenoid filter are currently only available for *Musca*^5,21^.

Our findings suggest that at the level of the photoreceptor, there is no detectable interaction between inner R7/R8 receptor pairs, or inhibition, which has been detected further downstream in the visual pathway at the first visual synapse in the medulla and *via* the Dm9 pathway^1,2^. This strongly suggests that there is no direct enhancement or inhibition of signal, which could be achieved by gap junctions between neighbouring photoreceptors or by local electrical fields in the surrounding extracellular space. Furthermore, although spectral inhibition occurs between photoreceptor terminals in the medulla^1,2^ it is not detectable upstream in *Drosophila*. If such inhibitory processes typically originate at the terminals and further downstream in other insects, then this may explain why so few studies have described spectral inhibition using intracellular recordings alone. Our results provide evidence for interactions between outer and inner receptors, which may indicate a feedback pathway in the lamina, where both inner and outer receptors interact. Cross-modulation *via* gap junctions in the lamina is known to occur between the Rh1-mediated motion pathway and R7/R8 colour pathway^22,23^ and was proposed as a mechanism to improve motion discrimination^13^. We suggest that our findings provide further evidence to support this circuit model. The addition of responses from all single rescue genotypes is in good agreement with wild type responses, demonstrating that overall, the voltage output of the photoreceptors sum linearly at the level of the retina (Supplementary Figure **S5**).

Our experiments show the importance of *in vivo* measurements for the full characterisation of visual systems that take into account the modulating effects of screening from distal receptors and ocular pigments. We found that the response of R8y receptor is strongly bathochromically shifted by both the presence of screening pigment and the blue-absorbing yellow filter. The latter also plays a role in refining the sensitivity of the Rh4 UV-sensitive visual pigment, which would likely enhance spectral discrimination. These findings contribute to the greater understanding of the *Drosophila* visual system and will assist in guiding future visual experiments and visual system modelling for which it is vital that the underlying photoreceptor sensitivity is known.

## Methods

### Animals

All *Drosophila melanogaster* stocks were maintained on 12/12 h light/dark light cycle at 22°C. Flies were reared on either yellow cornmeal or yeast-glucose food, for tests of carotenoid deprivation. Photoreceptor activity was selectively recovered by expression of phospholipase C (PLC) under opsin promotors against a *norpA* background, generating single opsin rescue flies. Opsin rescue flies were generated with wild-type screening pigment (red eye), (*w[+] norpA.CS; Rh-norpA)* and reduced screening pigment (orange eye), by incorporation of the *mini-white* gene, *w[-] norpA; Rh-norpA*. Single rescue flies were crossed to generate double opsin rescue genotypes. Rh6 opsin was ectopically expressed in the outer receptors under the control of the Rh1 promotor in a white eye mutant background, *w[-] norpA; Rh1-norpA Rh1-Rh6; ninaE*. Oregon R flies were used to test wild-type response and a *norpA* mutant with no rescue used as a negative control, *w[-] norpA;+;+* (Supplementary Figure **S6**).

### Light stimulus

Light from a 150W xenon arc lamp was coupled to a monochromator with either 1200 or 2400 line-ruled diffraction grating. Long-pass filters were placed in the light path to filter optical harmonics produced by the 1200 grating (< 250 nm, WG280, Schott) and 2400 grating (< 400 nm, GG435, Schott). Intensity of the light was controlled by altering the width of the input and exit slits of the monochromator. Peak wavelength was controlled by the grating angle, yielding a testing range between 315 – 550 nm or 450 – 700 nm for the 1200 and 2400 gratings, respectively. Wavelength and photon flux were calibrated at the point of the fly. All spectral measurements were made using spectrophotometers (Avantes AvaSpec 2048 Single Channel spectrometer and Ocean FX, OceanOptics) calibrated to a known light source for measurements of irradiance (DH2000, OceanOptics). Spectral acquisition was controlled using custom Matlab scripts (v2018a, Mathworks) and conversions to irradiance were carried out according to the manufacturer’s instructions. Irradiance in µWatt cm^-2^ was converted to photon flux in photons^-1^m^-2^s^-1^ according to the equation:

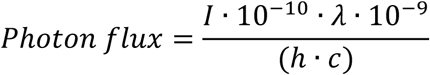

where *I* is irradiance,, *h* is Planck’s constant, *c* is the speed of light and *λ* is wavelength. A measure of total photon flux was calculated by integrating underneath the spectrum curve, which was used for all calibrations. To ensure isoquantal stimuli at each desired wavelength and intensity, we ran an automated calibration protocol using custom Matlab scripts (Data Acquisition Toolbox, Mathworks), which simultaneously controlled the spectrophotometer and monochromator. The position of entrance and exit slits of the monochromator and the diffraction grating were incrementally adjusted until the desired peak wavelength and total photon flux was reached. Each stimulus was calibrated to within +/- 0.5 nm peak wavelength, calculated using full width of spectrum at half maximum and to within +/- 0.75% total photon flux. All stimuli were then measured with the calibrated values to confirm calibration stability (Supplementary Figures **S7** and **S8**). Orange eye Rh1 and Rh5 rescue flies were illuminated with background light (670 nm) to increase recovery of sensitivity according to preliminary testing.

### ERG

Animals were anesthetised on ice and immobilised on a metal cone using ultraviolet curing adhesive (Norland). Electroretinogram (ERG) recordings were made using borosilicate micropipettes filled with insect saline. Recordings were measured from the equator of the eye and the reference electrode was positioned in the median ocellus. Light was delivered to the fly via a UV transmissive 5 mm liquid light guide and silica bi-convex lens with 25.4 mm focal length (Newport SBX019, USA), arranged to maximise light stimulation at the point of the fly. All recordings were made within a Faraday cage and responses were amplified (MultiClamp 700B amplifier, Molecular devices or EXT-02F, NPI). Both stimuli and data acquisition were controlled using a DAQ board (National Instruments) in conjunction with the software, Ephus^24^.

### Experimental design

To determine the response-log intensity (VlogI) function, each animal was tested with a series of 200 ms light pulses every 10 seconds that increased in intensity over a possible range of 6 log units. Red eye flies were tested between 1.14 × 10^7^ and 3.60 × 10^12^ photons^-1^m^-2^s^-1^. Orange eye flies were tested between 3.60 × 10^6^ and 6.40 × 10^11^ photons^-1^m^-2^s^-1^. Each intensity was repeated 10 times followed by a pause of 100 seconds. The wavelength each VlogI test was chosen according to previous estimates of peak sensitivity for the test visual pigment: Rh1, 485 nm; Rh3, 345 nm; Rh4, 370 nm; Rh5, 440 nm; Rh6 540 nm^4^. Peak wavelength for Rh6 was adjusted after preliminary tests indicated longer-wavelength peak sensitivity. The intensity at half maximum response was calculated from the VlogI curve and used for spectral tests. In cases where no obvious photoreceptor saturation had occurred, this value was estimated from the fitted curve.

To test spectral response, animals were stimulated with isoquantal flashes every 5 seconds at all test wavelengths with randomised presentation. Test wavelengths were divided into three blocks from lowest to highest wavelength and randomised within. Stimuli were always presented from these categories in order from low to high, to ensure a balanced order of testing across the wavelength range. Each wavelength was tested 10 times concurrently and the last 5 responses were used for analysis. All genotypes were tested with wavelengths of 315 – 550 nm and those with long wavelength responses (e.g. Rh6) were also tested with 450 – 700 nm, in steps of 5 nm. All animals were dark adapted for 30 minutes prior to the VlogI test and subsequently a further 15 minutes before each spectral sensitivity test.

### Analysis

ERG responses were normalised to a zero baseline using the average of 100 ms prior to stimulus onset. Photoreceptor response was calculated as the change in voltage between the zero baseline and minimum voltage during the 10 ms before the end of the light flash. The last 5 photoreceptor responses from each set of 10 repeats was used for VlogI and spectral sensitivity tests. These responses were averaged and mean responses used to compare genotypes. Animals with low or noisy ERG responses indicating a poor-quality preparation or inadequate electrode connection were not used for further analysis. VlogI data were fitted to the Naka-Rushton function:

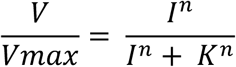

where *V* is the photoreceptor response, *Vmax* is the maximum response, I is the light intensity and *K* is the light intensity required to achieve half of *Vmax*^25^ and *n* is the slope. The intensity at half maximum response (K) was used for spectral tests. Spectral sensitivity data were first smoothed using a Savitzky-Golay filter (data window 15 nm) then averaged across the normalised data from all individuals in each experiment. To combine spectral sensitivity curves from tests using both the lower and higher wavelength gratings all non-normalised curves were joined at the 450 – 550 nm overlap region and an average fit was derived from the fit of all 21 points. The joined curves were then normalized between 0 and 1 and analysed as outlined previously. For a comparison of photoreceptor pair responses, the mean of non-normalised sensitivity curves from single rescue flies were summed pair-wise according to the order of light stimulus and compared with the response curves of double rescue flies. All analyses were performed in MATLAB (v2018b, Mathworks), using custom scripts. Visual pigment templates were generated using R package PAVO ^26^. Two sample Student’s t-tests were carried out with Bonferroni correction for multiple sampling between sensitivity curves with the exception of comparisons made between double opsin rescues tested at different light intensities, which were tested using paired t-tests. All statistical tests were carried out in R^27^.

## Acknowledgements

We thank Cairn Research (in particular Jeremy Graham) for assistance with customisation of the optoscan monochromator and the University of Cambridge fabrication workshops. We also thank Ilse Daly and Nicholas Roberts for their assistance with spectrophotometer calibrations.

## Author contributions

CRS collected, analysed, interpreted data and wrote the manuscript. JB generated transgenic flies. MML and DPB collected ERG data. TW conceived the study and assisted with data interpretation. CRS, JB and TW edited the manuscript.

## Funding

We thank the Biotechnology and Biological Sciences Research Council (BBSRC) for a David Phillips Fellowship to support TJW (BB/L024667/1) and the College of Biological Sciences, University of Minnesota.

## Competing interests

The authors declare no competing interests.

