## Supplementary Data for "The spectral sensitivity of *Drosophila* photoreceptors"

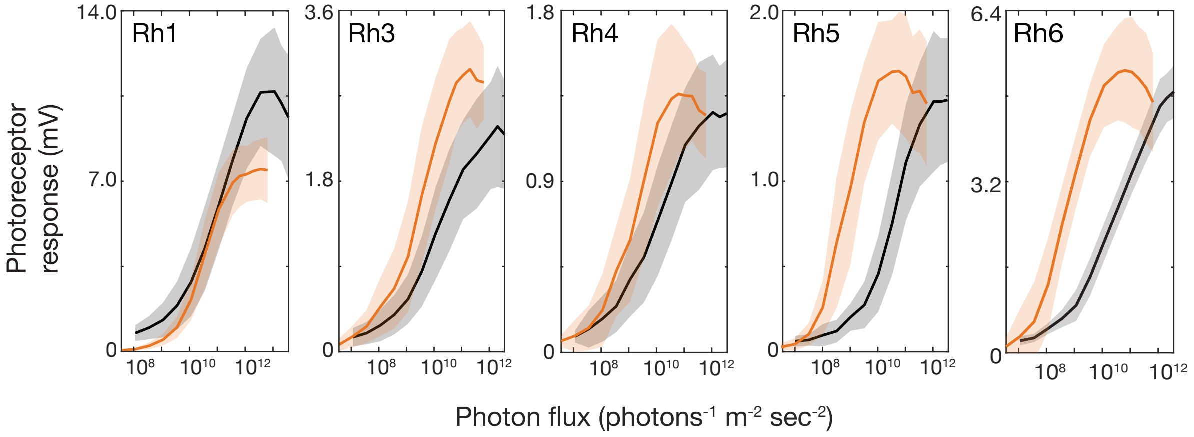


**S1.** Voltage response of photoreceptors in single rescue flies (Rh1, Rh3 - 6) with red (black lines) or low levels of screening pigment (orange lines), tested over 6 log units of light. Photon flux is on a logarithmic scale. Error shown is standard deviation.

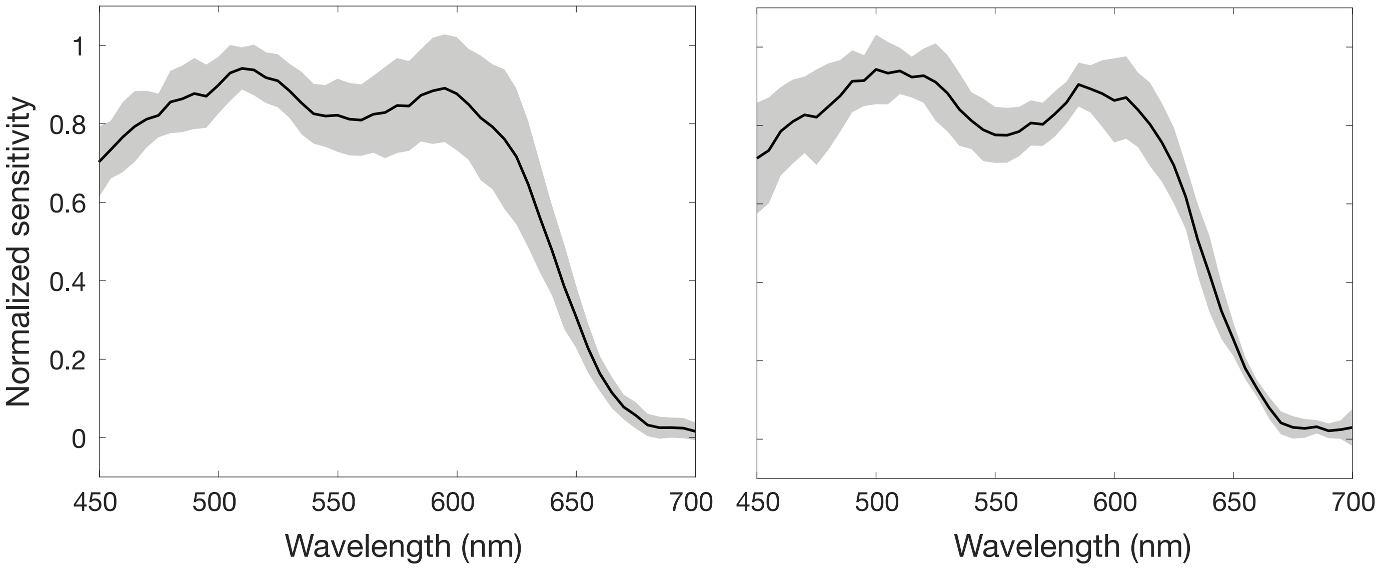


**S2.** Spectral response of red-eye Rh6 rescue flies with one (left) or two (right) generations of carotenoid deprivation. Error shown is standard deviation.

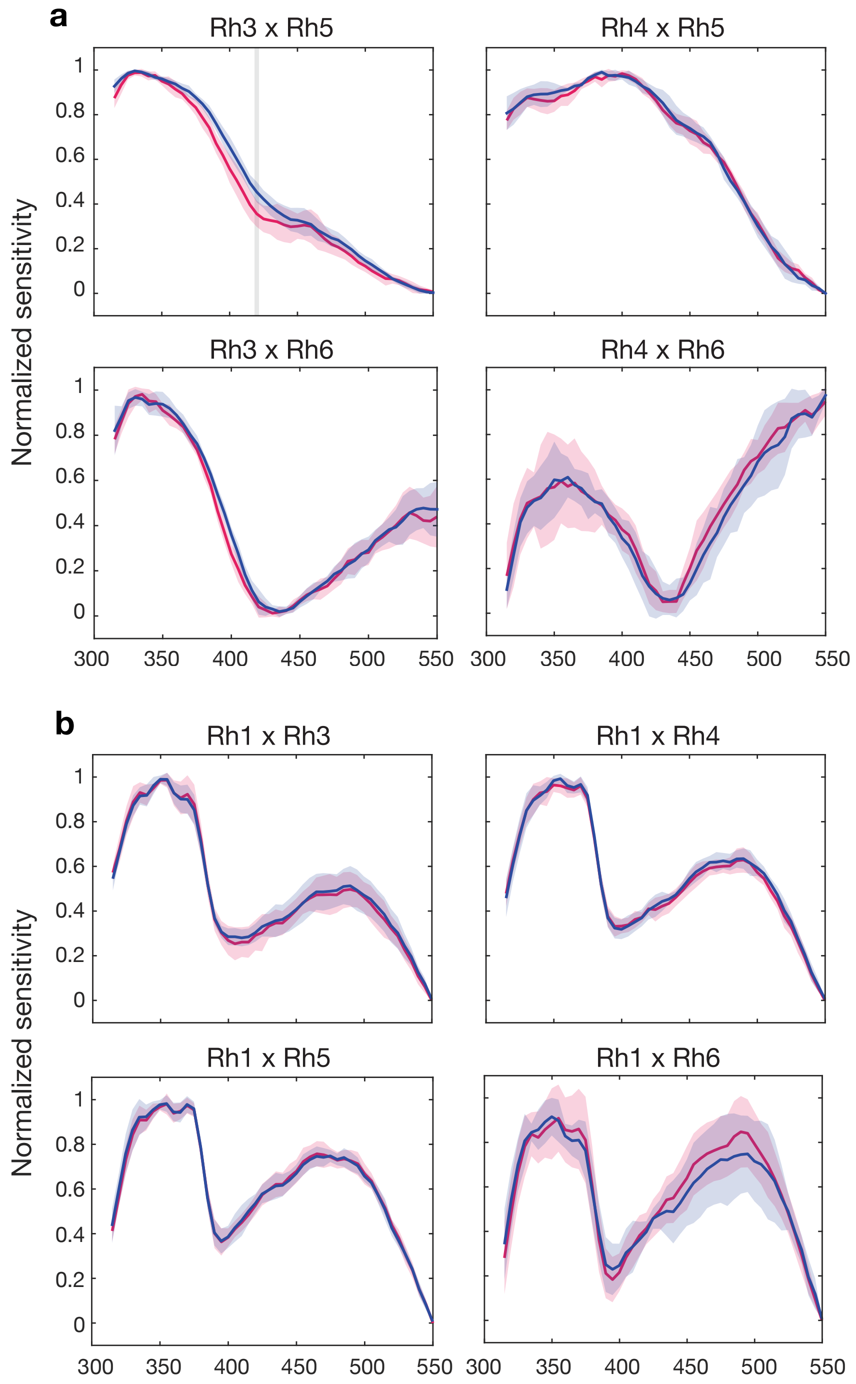


**S3.** (a) Spectral sensitivity curves of double opsin rescues with two inner photoreceptor types active. Double opsin rescue flies were tested at an intensity derived from the V-logI test at either the Rh3/Rh5 peak sensitivity (pink) or Rh5/Rh6 peak sensitivity (blue). A significant change in shape was only observed in Rh3 x Rh5 flies at 420 nm. (b) Spectral sensitivity curves of double opsin rescues with one inner photoreceptor type and outer receptors active. Flies were tested at an intensity derived from the V-logI test at either the Rh1 peak sensitivity (pink) or at the peak sensitivty of the inner receptor opsin Rh3 - Rh6 (blue). Shading denotes significance between sensitivity curves using a paired Student's t-test at $p<0.001$. Error shown is standard deviation.

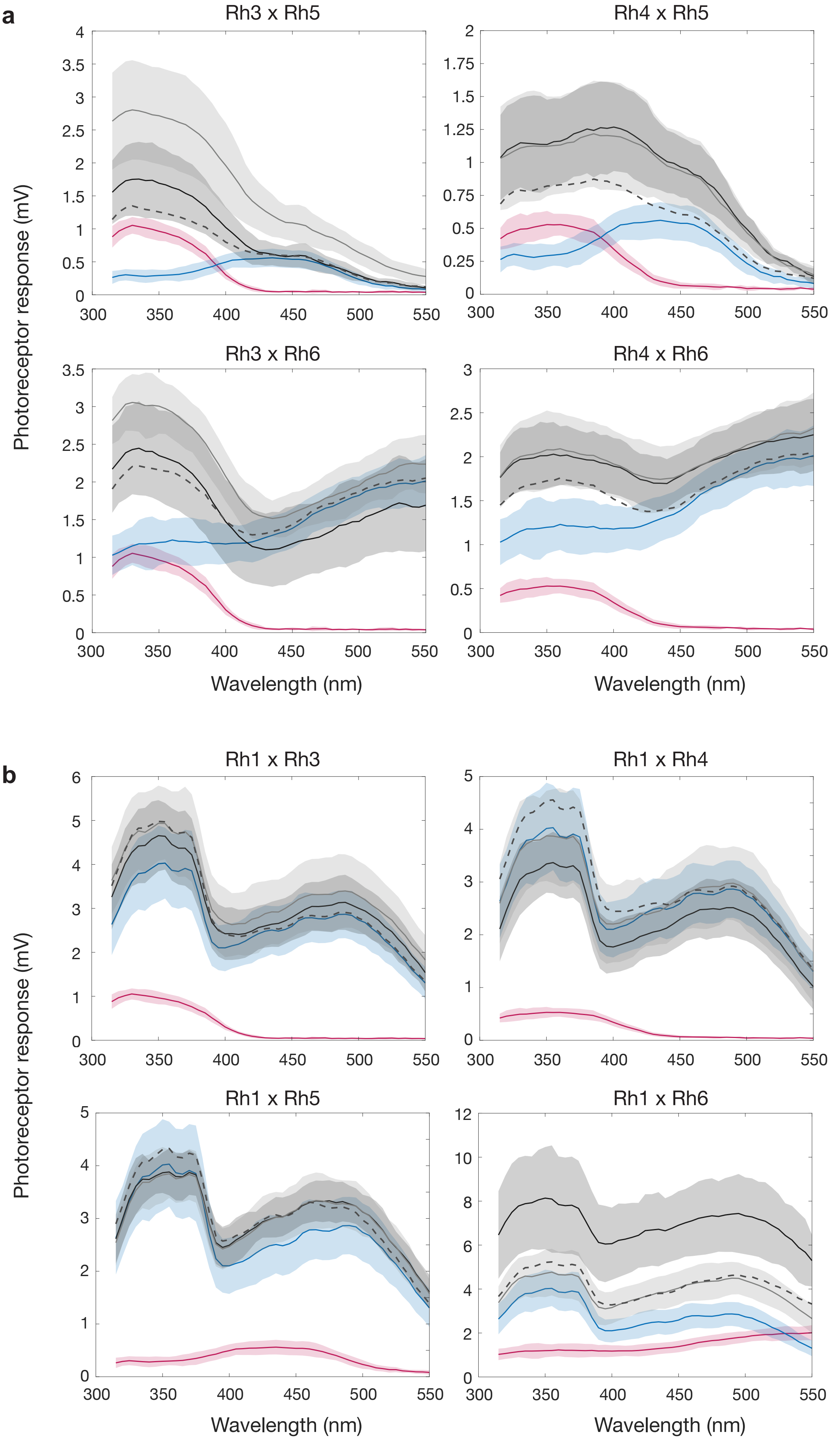


**S4.** (a) Voltage responses of single (pink: Rh3 or Rh4 and blue: Rh5 or Rh6) and double opsin rescue flies tested at an intensity derived from the VlogI response of either Rh3/Rh4 (dark grey) or Rh5/Rh6 (light grey). The algebraic sum of the single responses (dashed lines). (b) Voltage responses of single (pink: Rh3 - Rh6 or blue: Rh1) and double opsin rescue flies tested at an intensity derived from the VlogI response of either Rh3 - Rh6 (dark grey) or Rh1 (light grey). Error shown is standard deviation.

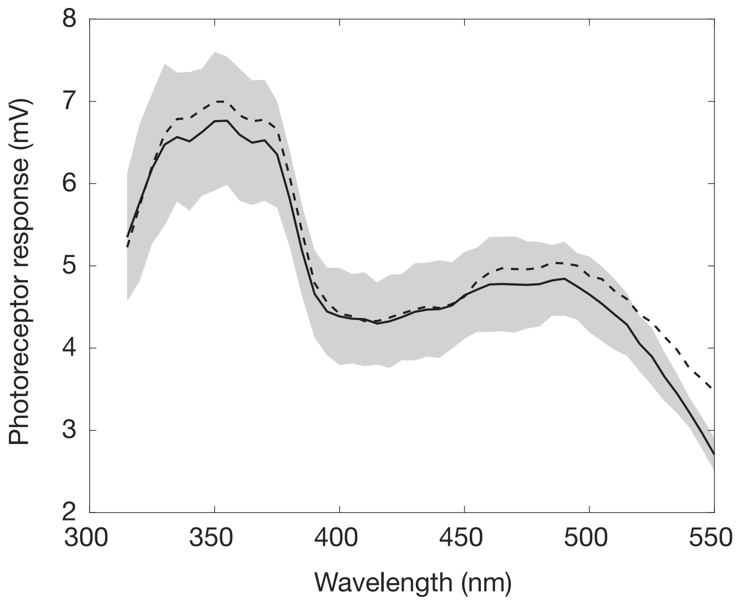


**S5.** Photoreceptor response of wild-type flies (solid line) and the algebraic sum of mean responses from all red-eye single opsin rescues (dashed line). Error shown is standard deviation.

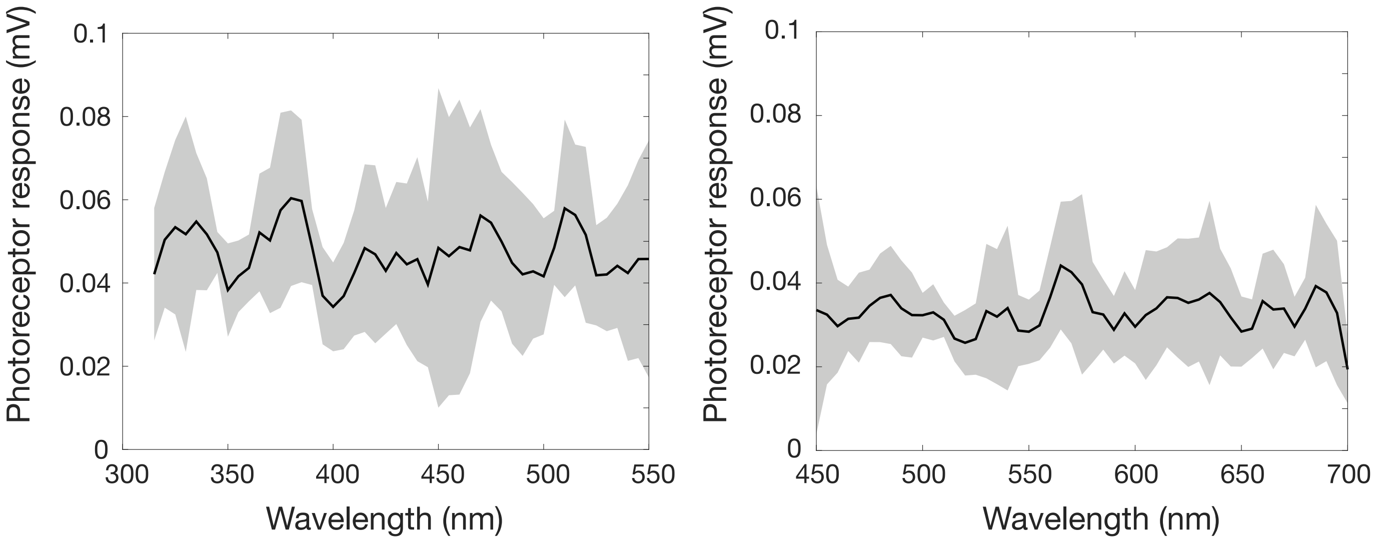


**S6.** Photoreceptor response of control flies *w[-] norpA;+;+* indicating the baseline response from flies with no photoreceptor response. Control flies were tested across the test wavelength ranges: 315 - 550 nm and 450 - 700 nm. Flies were tested at the maximum possible intensity. Error shown is standard deviation.

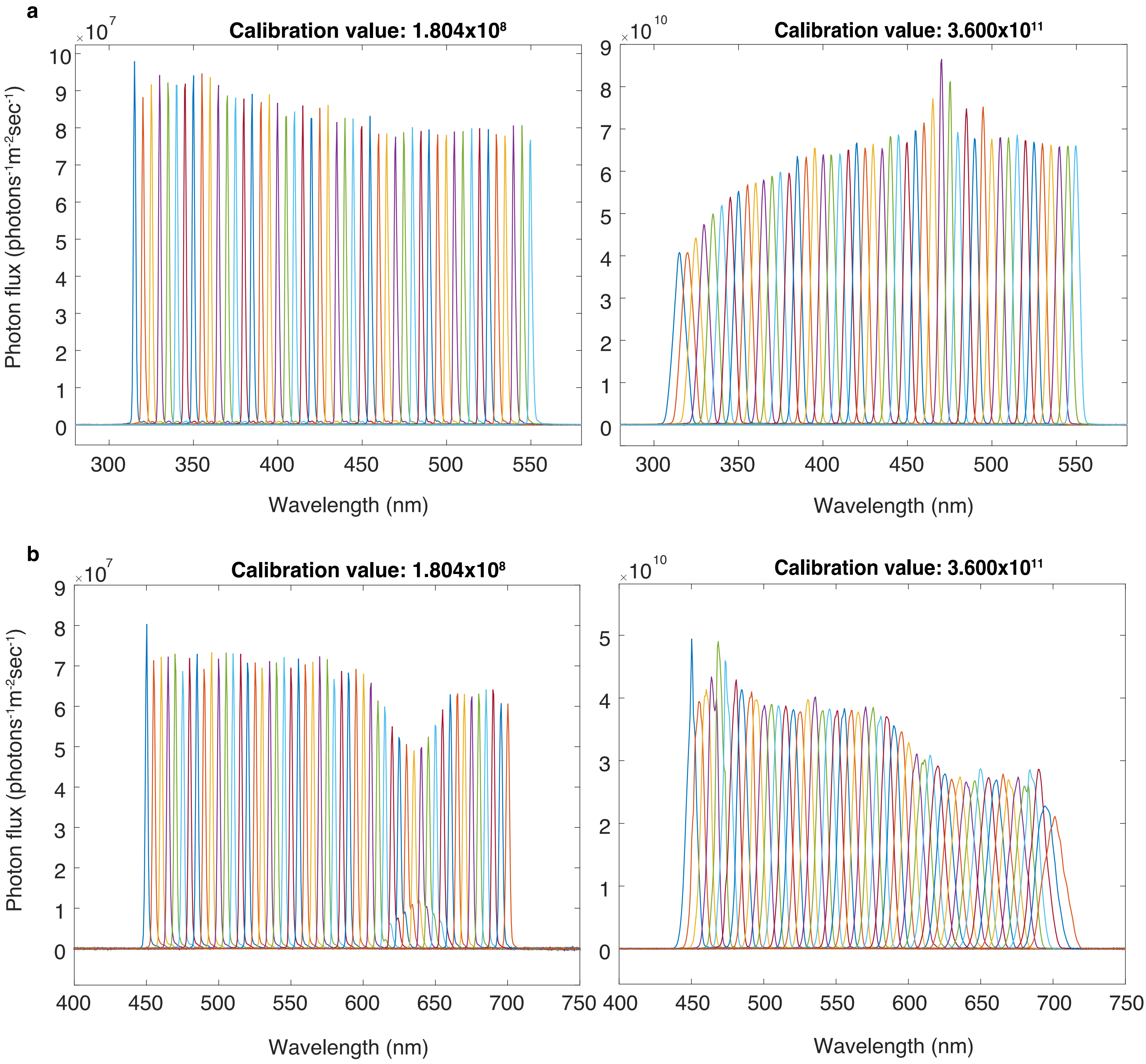


**S7.** Example spectra of isoquantal stimuli measured after calibration to low ($1.804 \times 10^8$) and high ($3.6 \times 10^{11}$) target calibration values of total photon flux (area under the spectrum). Each spectrum represents one calibration point, in 5 nm steps from 315 - 550 nm (a) and 450 - 700 nm (b).

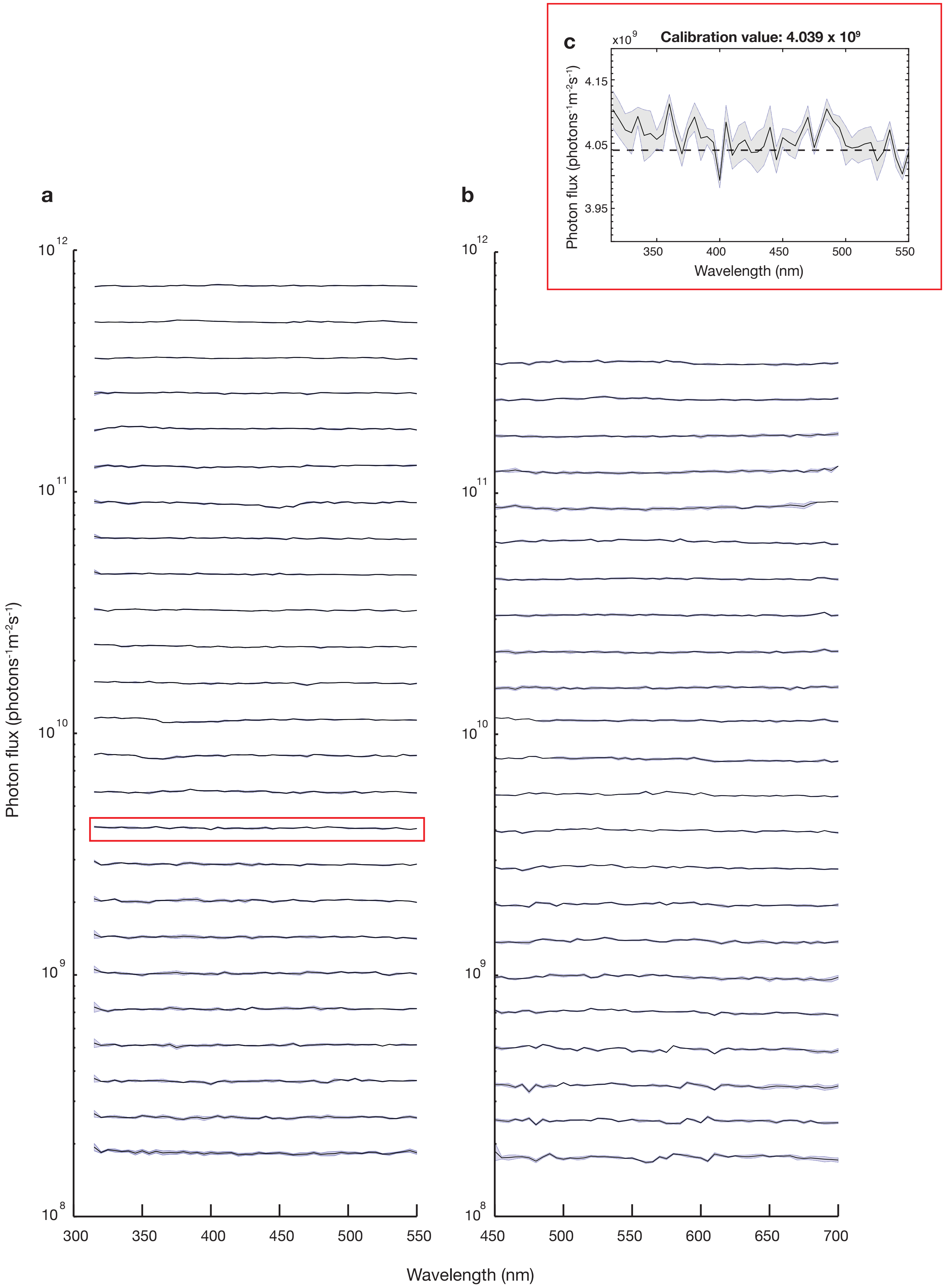


**S8.** Example of measurements made after calibration to spectral test intensities and wavelengths for (a) the 1200 and (b) 2400 line ruled diffraction grating. The y axis is on a log scale. Five measurements were taken per calibration point and the average taken. Magnified version of indicated calibration measurements (c) with calibration reference vaue shown (dashed line). Error shown is standard deviation.
